# Deficiency of the lysosomal protein CLN5 alters lysosomal function and movement

**DOI:** 10.1101/2021.08.24.457390

**Authors:** Indranil Basak, Rachel A. Hansen, Michael E. Ward, Stephanie M. Hughes

## Abstract

Batten disease is a devastating childhood rare neurodegenerative disease characterized by rapid deterioration of cognition and movement, leading to death within ten to thirty years of age. One of the thirteen Batten disease forms, *CLN5* Batten disease, is caused by mutations in the *CLN5* gene leading to motor deficits, mental deterioration, cognitive impairment, visual impairment, and epileptic seizures in children. A characteristic pathology in *CLN5* Batten disease is the defects in lysosomes, leading to neuronal dysfunction. In this study, we aimed to investigate the lysosomal changes in CLN5-deficient human neurons. We used an induced pluripotent stem cell system, which generates pure human cortical-like glutamatergic neurons. Using CRISPRi, we inhibited the expression of *CLN5* in human neurons. The CLN5-deficient human neurons showed neutralised lysosomal acidity and reduced lysosomal enzyme activity measured by microscopy and flow cytometry. Furthermore, the CLN5-deficient human neurons also showed impaired lysosomal movement – a phenotype that has never been reported in *CLN5* Batten disease. Lysosomal trafficking is key to maintain local degradation of cellular wastes, especially in long neuronal projections and our results from the human neuronal model present a key finding to understand the underlying lysosomal pathology in neurodegenerative diseases.

## 1. Introduction

Batten disease or neuronal ceroid lipofuscinosis (NCL) is a group of rare, fatal inherited neurological diseases predominantly affecting children and is characterized by symptoms including motor deficits, mental retardation, cognitive impairment, visual impairment, and epileptic seizures (reviewed in [1]). There are thirteen forms of Batten disease caused by mutations in thirteen *CLN* genes (*CLN1-8* and *10-14*)[2, 3]. Mutations in one of the *CLN* genes, *CLN5*, cause a variant late-infantile NCL[4–6]. Life expectancy for children with *CLN5* mutations is between ten and thirty years, and there is currently no cure or approved treatment for CLN5 Batten disease[1, 4, 6, 7].

CLN5 is a lysosomal protein with ambiguous protein function (reviewed in [1]). A cellular hallmark of Batten disease is defective lysosomes, the cellular waste recycling machinery[3]. Lysosomes are acidic vesicles containing hydrolytic enzymes, and loss of CLN5 shows reduced acidic organelles and impaired autophagy, as observed in ovine neural cultures from naturally occurring CLN5-deficient sheep model [8]. To understand what other properties of the lysosomes are impaired due to the loss of CLN5 leading to neuronal dysfunction, we inhibited CLN5 expression in human neurons. In this study, we developed a unique CLN5-deficient human neuronal model with isogenic controls from induced pluripotent stem cells (iPSCs) using clustered regularly interspaced short palindromic repeat interference (CRISPRi). In our CLN5-deficient human neuronal model, we confirm the reduced acidic organelles, as observed in our previous ovine study[8]. Intriguingly, we report, for the first time, that CLN5 deficiency in human neurons show impaired lysosomal enzyme activity and impaired lysosomal movement, both being crucial for lysosomal homeostasis and hence, maintaining neuronal health.

## 2. Materials and Methods

### 2.1. Generation of induced pluripotent stem cell derived human neurons

For this study, we used a published protocol established by Dr. Michael Ward’s group[9] to generate pure human cortical-like glutamatergic neurons from iPSCs. In short, a transcription factor *neurogenin-2* in a doxycycline inducible cassette was stably integrated into the AAVS1 safe harbour locus of iPSCs cultured in Essential 8 media (ThermoFisher Scientific, A1517001). The expression of *neurogenin-2* in the iPSCs was induced for three days using doxycycline (Sigma-Aldrich, D9891). Post-induction, the differentiation of the *neurogenin-2* iPSCs into mature isogenic, integrated and inducible pure human cortical-like glutamatergic neurons (i^3^Ns) was completed in two weeks in BrainPhys media (STEMCELL Technologies, 05790) (Figure. 1A). After two weeks, the iPSC-derived mature neurons were fixed with 4 % paraformaldehyde (Sigma-Aldrich, P6148). The neurons were recognized using antibodies for neuronal markers (listed in Table S1) and stained with 4’, 6-diamidino-2-phenylindole (DAPI) (Sigma-Aldrich, D9542), as previously described[8]. The stained neurons were imaged (Figure 1B) on a Nikon A1R resonant scanner confocal microscope.

**Figure 1.**
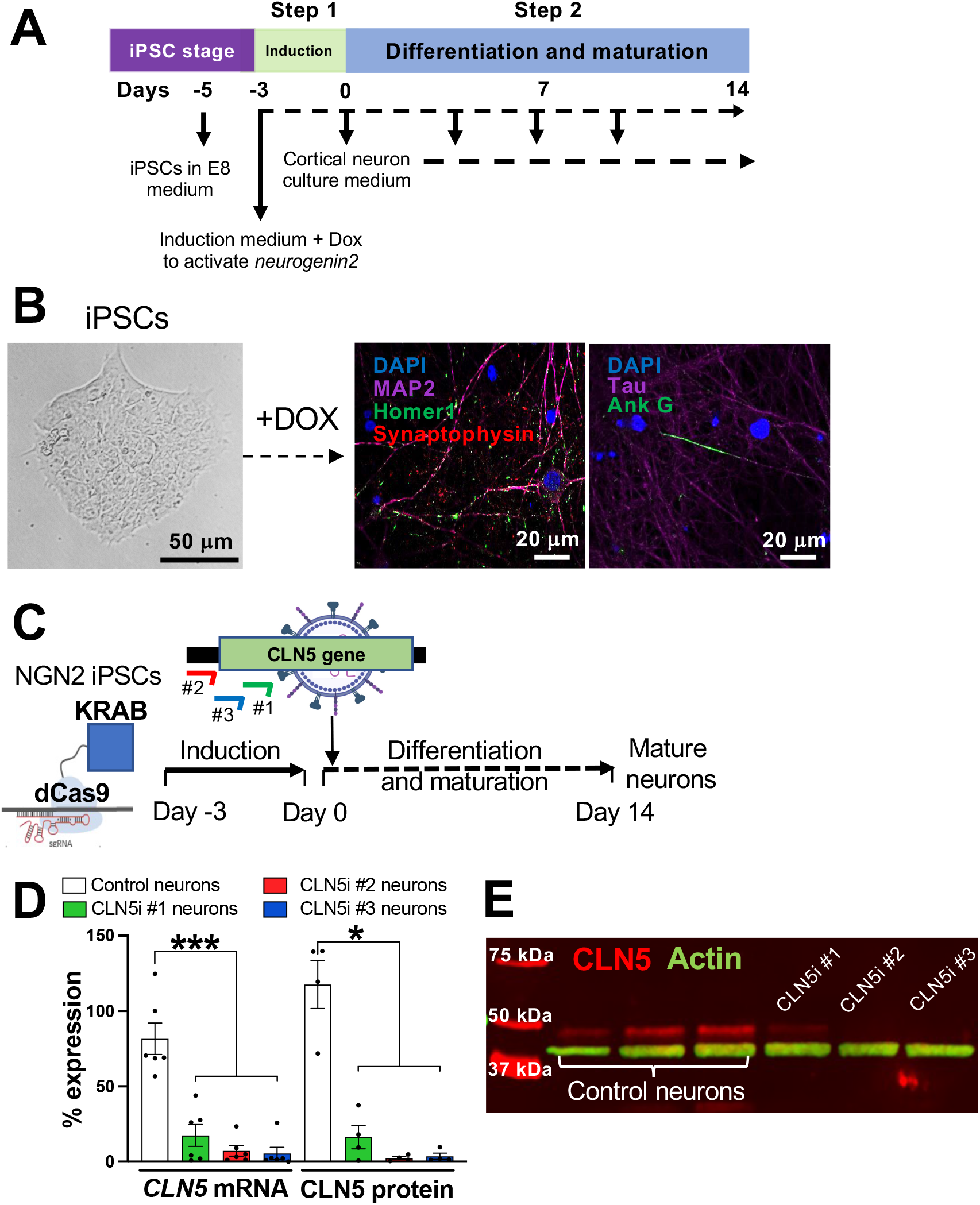
iPSC-derived human cortical-like glutamatergic neurons mimicking CLN5 Batten disease. (A) Schematic of iPSC-derived human neuron generation. Three days of induction and fourteen days of differentiation and maturation gives pure human cortical-like glutamatergic neurons (B) iPSCs with integrated *neurogenin-2* can be efficiently differentiated into pure human cortical-like glutamatergic neurons expressing neuronal markers. (C) CRISPRi strategy to inhibit *CLN5* in iPSC-derived human neurons. Three sgRNAs against *CLN5* were transduced using lentiviruses into iPSC-derived human neurons containing dCas9 machinery. (D, E) All three sgRNAs showed significant inhibition of CLN5, both at the transcript (D) and protein level (D, E). (E) First three lanes are control neurons, and next three lanes are CLN5 neurons. n ≥ 4, * p < 0.05, *** p < 0.001.

### 2.2 CRISPR interference in iPSC-derived neurons

*Neurogenin-2* iPSCs with deactivated Cas9 (dCas9-KRAB)[10] tagged with a blue fluorescent protein (BFP) stably integrated with into the CLYBL safe harbour locus[9] were induced to generate i^3^Ns, which have been used for the following experiments from at least three different passage numbers. To inhibit the expression of *CLN5*, three individual single guide RNAs (sgRNAs) were designed against CLN5 – two in the coding region (#1 and #3) and one in CLN5 5’UTR (#2) (Figure 1C), following Horlbeck *et al*. [11]. A control sgRNA was designed against green fluorescent protein[12]. Following previously established protocols [2], the control and CLN5 sgRNAs were cloned into Addgene plasmid #60955 (pU6-sgRNA EF1Alpha-puro-T2A-BFP) and then packaged into lentiviruses (Addgene plasmid #12259 and 12260). To inhibit CLN5 in dCas9-i^3^Ns (henceforth termed as CLN5i neurons), induced dCas9-i^3^Ns were transduced with lentiviruses with either CLN5 sgRNAs or control sgRNA on Day 1 of differentiation / maturation (Figure 1C) at a multiplicity of infection (MOI) of 1. The lentivirally transduced neurons were maintained until Day 14 with half media changes on Days 4, 7 and 10 (Figure 1A). The Day 14 control and CLN5i neurons were harvested with Accutase (ThermoFisher Scientific, A1110501) to isolate total RNA using PureLink RNA isolation kit (ThermoFisher Scientific, 12183018A), following manufacturer’s instructions. The RNA isolates were treated with DNAseI (ThermoFisher Scientific, 18-068-015) and quantified using a Nanodrop spectrophotometer, following our previously established protocol[8]. *CLN5* transcripts were measured from the control and CLN5i neurons RNA by quantitative polymerase chain reaction[13] using an iTaq Universal Probes one-step kit (Bio-Rad, 1725141) on a Roche 96-well thermocycler, following manufacturer’s instruction. Similarly, Day 14 control and CLN5i neurons were used to isolate total cell lysates using N-PER Neuronal Protein Extraction Reagent (ThermoFisher Scientific, 87792), following manufacturer’s instructions. The protein isolates were quantified and tested for CLN5 protein expression using western blot, as previously described[14]. *GAPDH* [13] and ACTIN were used for normalisation for the qPCR and western blot analyses, respectively. Details of the qPCR probes and antibodies used are listed in Table S1.

### 2.3 Assessment of lysosomal acidity, number, enzyme activity and movement

To test the lysosomal acidity in control and CLN5i neurons, LysoTracker Red DND 99 dye (ThermoFisher Scientific, L7528) was used following our previously published protocol[8]. Induced control and CLN5i neurons were cultured for two weeks in BrainPhys media on 96-well clear bottom plate (Corning, 3599) (plated at 10,000 neurons/well for high-throughput image analysis), on Nunc Lab-Tek II Chamber Coverglass (ThermoFisher Scientific, 155409PK) (plated at 30,000 neurons/well for confocal microscopy), and on 24-well cell culture plate (Corning, 3524) (plated at 100,000 neurons/well for Flow Cytometry). The Day 14 mature neurons were stained with the LysoTracker Red dye following Best *et al*.[8] The LysoTracker Red stained control and CLN5i neurons were used for obtaining still images as well as videos using an Olympus FV3000 confocal microscope. For high-content image analysis, the LysoTracker Red stained neurons were further stained with Hoechst stain for 10 minutes at 37 °C, maintained in Phenol Red-free BrainPhys media (STEMCELL Technologies, 05791), imaged on a Cytation 5 cell-imaging multi-mode reader (BioTek) and analysed using Fiji/ImageJ[15]. For the flow cytometric analysis, neurons were stained as above followed by incubation with Accutase at 37 °C for 5 minutes. After Accutase incubation neurons were harvested and resuspended in Phenol Red-free BrainPhys media followed by recording LysoTracker Red mean fluorescence intensity on a BD Fortessa (BD Biosciences).

To test the change in the number of lysosomes, we transduced the control and CLN5i neurons with a lentivirus expressing lysosome-associated membrane protein 1 (LAMP1) tagged with neonGreen that would target all lysosomes and early endosomes. The neonGreen tagged organelles in control and CLN5i neurons were quantified using flow cytometry, as described above.

For assessing cathepsin B activity (Magic Red assay kit, Abcam, 270772) in the control and CLN5i neurons, Magic Red dye was diluted 1:150 and 1:250 for high-content imaging and confocal imaging, respectively, in Phenol Red-free BrainPhys media. The control and CLN5i neurons were incubated with the diluted Magic Red dye for 30 minutes at 37°C followed by staining with Hoechst stain for 10 minutes at 37 °C. The stained neurons were maintained in Phenol Red-free BrainPhys media for imaging as described earlier.

For the lysosomal movement analysis, time-lapse videos were obtained on an Olympus FV3000 confocal microscope. For one video, 60 images were taken at 1 frame per 5 second for 5 minutes. At least two videos were obtained per sample, with ten kymographs generated per video and the experiments were performed in three independent replicates. The resulting videos were analysed using Kymograph Clear[16] to generate ten kymographs per video. The kymographs were further analysed for lysosomal movement using KymoButler[17].

### 2.4 Statistical analysis

All experiments were performed n ≥ 3 times and analysed on GraphPad Prism (GraphPad). For RNA, protein, Lysotracker Red microscopic and flow cytometry analyses, and Magic Red microscopic analysis, one-way ANOVA was used to assess statistical significance. For the lysosomal movement analyses, in addition to one-way, we performed a two-way ANOVA analysis the overall change in lysosomal movement in CLN5i neurons compared to control neurons. Data has been presented as Mean ± standard error of mean (SEM). * p < 0.05, ** p < 0.01, *** p < 0.001.

## 3. Results

### 3.1. Inhibition of CLN5 in human neurons

In our iPSC-derived neurons (Figure 1A), we confirmed the expression of neuronal markers (Figure 1B) – MAP2 for dendrites, Tau for axons, Synaptophysin for pre-synaptic vesicles, Homer1 for postsynaptic structures and Ankyrin G for axon initial segment. The CRISPRi strategy adapted to inhibit the expression of *CLN5* in neurons gave more than 90 % CLN5 inhibition for two sgRNAs (CLN5i #2 and #3 neurons) compared to the control, both at the transcript level (Figure 1D) and at the protein level (Figure 1D, E). The expression of *CLN5* in CLN5i #1 neurons was more than CLN5i #2 and #3 neurons, albeit lower than the control neurons. Hence, we continued using all three sgRNAs for the following analyses where CLN5i #2 and #3 neurons represent homozygous CLN5 knockout neurons and CLN5i #1 neurons represent heterozygous CLN5 neurons. Our CLN5-deficient neurons are the first human neuronal model with isogenic controls to study the associated neuronal pathogenesis.

### 3.2. Loss of CLN5 impairs lysosomal acidity and lysosomal enzyme activity in human neurons

Like our previously published experiments in CLN5 ovine cells, we observed reduced acidic organelles, without a change in total number of lysosomes[8] in our iPSC-derived human neuronal model (Figure 2A-E). LysoTracker Red stains acidic lysosomes, and the CLN5i #2 and #3 neurons showed significant decrease in Lysotracker Red signal compared to control neurons (Figure 2A, B) (% LysoTracker Red fluorescence/neuron: Mean±SEM for Control vs CLN5i #1, #2, #3 neurons: 100 vs 93.76±8.99, 78.76±3.89, 80.61±2.98; p value Control vs CLN5i #1, #2, #3 neurons: 0.801, 0.013, 0.007). We verified our LysoTracker Red microscopy results by flow cytometric analyses, which supported the decrease in Lysotracker Red signal in CLN5i neurons compared to control neurons (Figs. 2C, D) (LysoTracker Red mean fluorescence intensity/neuron: Mean±SEM for Control vs CLN5i #1, #2, #3 neurons: 100 vs 108±9.54, 59.94±3.61, 49.69±2.33, p value Control vs CLN5i neurons #1, #2, #3: 0.752, 0.0009, <0.0001). The total number of lysosomes, measured by quantifying LAMP1-neonGreen positive organelles did not show any significant difference between control and CLN5i neurons (Figure 2E) (LAMP1-neonGreen median fluorescence intensity/neuron: Median±SEM for Control vs CLN5i #1, #2, #3 neurons: 100 vs 96.38±11.38, 104.7±18.11, 107±17.04, p value Control vs CLN5i neurons #1, #2, #3: 0.977, 0.987, 0.953). Our result indicates that the number of lysosomes in the neurons are not altered by inhibiting CLN5, however, lysosomal acidity is neutralised when CLN5 is inhibited.

**Figure 2.**
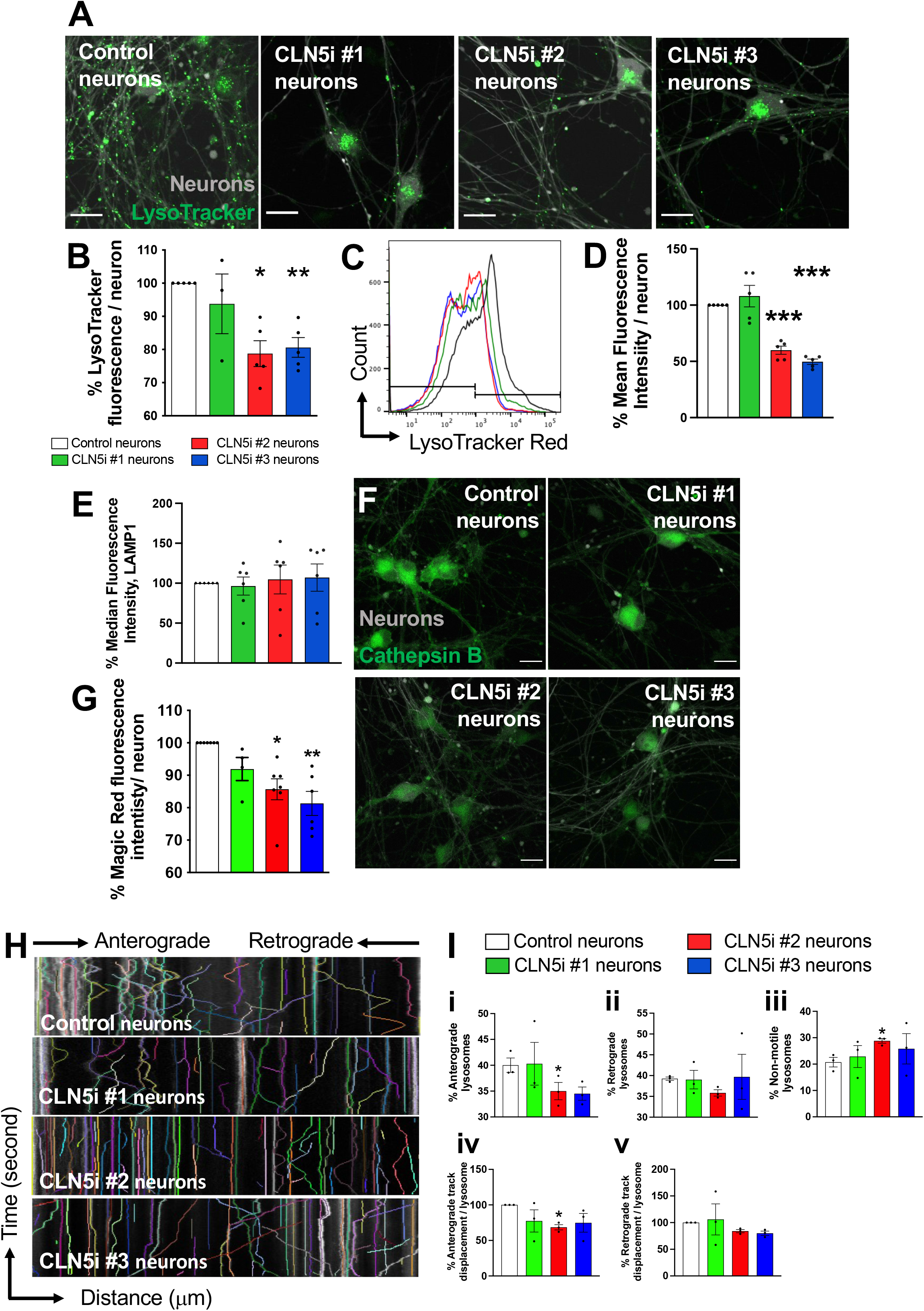
CLN5-deficient human neurons exhibit defects in lysosomal acidity, enzyme activity and movement. (A, B) Microscopic analysis of lysosomal acidity in the iPSC-derived human neurons revealed neutralised lysosomal acidity (measured by LysoTracker, compared pseudo colour green dots) in CLN5i neurons (#2 and #3) compared to control neurons. (C, D) The microscopic analysis of lysosomal acidity was validated by flow cytometric analysis. (E) The CLN5i human neurons did not show difference in lysosome number, measured by quantification of LAMP1-neuonGreen positive organelles using flow cytometry. (F, G) Microscopic analysis of lysosomal cathepsin B activity in the iPSC-derived human neurons revealed reduced enzyme activity (measured by Magic Red assay, compare pseudo colour green) in CLN5-deficient neurons (#2 and #3) compared to control neurons. (H, I) CLN5-deficient neurons (CLN5i #2 neurons) showed reduced percentage of anterograde lysosomes, reduced anterograde track displacement and increased percentage of non-motile neurons compared to control neurons measured by confocal microscopy and kymograph analysis. n ≥ 3, * p < 0.05, ** p < 0.01, *** p < 0.001. Scale bar 20 μm.

Like the lysosomal acidity phenotype, the Magic Red stained control and CLN5i Day 14 neurons (Figure 2F, G), showed a decrease in cathepsin B activity (% Magic Red fluorescence intensity/neuron: Mean±SEM for Control vs CLN5i #1, #2, #3 neurons: 100 vs 91.9±3.57, 85.68±3.22, 81.31±3.69, p value Control vs CLN5i neurons #1, #2, #3: 0.173, 0.011, 0.008). Our results suggest that the neutralisation of lysosomal acidity is affecting the activity of the lysosomal cathepsin B in the CLN5i neurons compared to the control neurons.

### 3.3. Loss of CLN5 impairs lysosomal movement in human neurons

Lysosomes travel to distal ends of neurons to maintain local degradative activities, therefore making lysosomal movement crucial to maintain homeostasis in the neurons[18]. Some CLN proteins have been recently shown to alter lysosomal positioning[19, 20], however whether inhibiting CLN5 changes the lysosomal movement/trafficking in human neurons is unknown. To assess the lysosomal movement in control vs CLN5i neurons, kymographs were generated from time lapse videos of LysoTracker Red stained lysosomes in control vs CLN5i neurons (Figure 2H). The kymographs generated show three features – 1) curved lines going from left to right represent lysosomes moving in the anterograde direction (i.e., away from the cell body), 2) curved lines going from right to left represent lysosomes moving in the retrograde direction (i.e., towards the cell body), and 3) straight lines representing non-motile lysosomes. The CLN5i neurons, especially CLN5i #2 neurons (Figure 2Ii), showed significantly reduced percentage of anterograde lysosomes, whereas CLN5i #3 neurons showed a similar trend (% anterograde lysosome: Mean±SEM for Control vs CLN5i #1, #2, #3 neurons: 40.02±1.39 vs 40.29±4.13, 35±1.67, 34.5±1.28, p value Control vs CLN5i neurons #1, #2, #3: 0.999, 0.041, 0.201) (Figure 2Ii). No statistically significant difference was observed for percentage of retrograde lysosomes, although CLN5i #2 neurons showed a decreasing trend compared to the control neurons (Figure 2Iii). CLN5i neurons showed an increased percentage of non-motile lysosomes compared to the control neurons, with CLN5i #2 neurons showing a significant increase (% non-motile lysosomes: Mean±SEM for Control vs CLN5i #1, #2, #3 neurons: 20.72±1.81 vs 22.87±4.16, 28.81±0.887 vs 25.8±5.76, p value Control vs CLN5i neurons #1, #2, #3: 0.731, 0.028, 0.561) (Figure 2Iiii). The direction of travel of the lysosomes, anterograde, retrograde, or non-motile, are dependable variables for lysosomal movement. To test whether the overall movement of lysosomes was impaired in CLN5i vs control neurons, a two-way ANOVA test was performed that showed significant difference in lysosomal movement between CLN5i vs control neurons (p < 0.01). When we tested the distance travelled by the lysosomes in control and CLN5i neurons, we observed that CLN5i #2 neurons showed a significant reduction in distance covered by the anterograde lysosomes (% anterograde lysosomal displacement per lysosome: Mean±SEM for Control vs CLN5i #1, #2, #3 neurons: 100 vs 77.40±15.63, 68.50±3.70, 74.79±13.13, p value Control vs CLN5i neurons #1, #2, #3: 0.494, 0.026, 0.351) (Figure 2Iiv). CLN5i #2 and #3 neurons showed a decreasing trend in distance covered by the retrograde lysosomes compared to the control neurons, however, no statistical significance was achieved (Figure 2Iv).

Interestingly, we also observed some lysosomes that were motile but could not decide on the direction of their movement. As a result, these bidirectionally motile lysosomes kept moving back and forth without actually moving in their designated direction. This phenotype was prominent in CLN5i #2 and #3 neurons (Video S1B, C) as compared to control neurons (Video S1A), which showed smooth uni-directional movement of the lysosomes.

## 4. Discussion

Our study, involving a unique human neuronal model mimicking CLN5 Batten disease with isogenic controls, revealed - 1) our capability to model Batten disease in a dish using CRISPRi and iPSC-derived human neurons 2) > 90 % inhibition of CLN5 expression was achieved using CRISPRi in iPSC-derived human neurons, and CLN5-deficient human neurons manifest defects in 3) lysosomal acidity, 4) lysosomal enzyme activity, and 5) lysosomal movement. Our study is the first one to report that CLN5 deficiency in human neurons can lead to impaired lysosomal enzyme activity and lysosomal movement.

Lysosomal acidity is crucial to maintain a working environment for the lysosomal degradative enzymes. Hence, neutralisation of lysosomal acidity (Figure 2A-D) could render the enzymes inactive (Figure 2F, G), which, in turn could impair degradation of cellular waste, ultimately leading to toxic buildup of cellular waste and increase in neurotoxicity. It will be interesting to test other lysosome enzyme activities and whether their total protein levels change with the inhibition of CLN5 in the human neurons.

Lysosomes travel throughout the neurons, including the axons and the dendrites, as lysosomal transport is crucial not only for waste clearance in the neuronal projections, but also for the lysosomal secondary neuronal functions like synapse regulation and cargo transport[21–24]. Retrograde transport allows the lysosomes to fuse with autophagosomes near the cell body in the neurons [25], whereas anterograde lysosomes are known to carry on degradative activities at the distal ends of the axons[18]. In our *in vitro* human neuronal cultures, to include both axonal and dendritic functional anterograde and retrograde lysosomes, we imaged the LysoTracker positive acidic lysosomes to understand the effect of CLN5 inhibition on lysosomal movement in the whole neuron. From our data, it is apparent that anterograde lysosomal movement is impaired, whereas retrograde lysosomal movement appears to be impaired in CLN5-deficient human neurons. Dysfunctional lysosomal trafficking could result in the failure of the lysosomes to fuse with the autophagosomes, leading to an impaired autophagy-lysosomal pathway, as seen in our previous study[8]. Impaired lysosomal function in neurodegenerative diseases is not uncommon[26, 27]. Hence, it will be interesting to test whether other forms of Batten disease as well as other neurodegenerative diseases show such lysosomal trafficking anomalies. Furthermore, testing whether the expression or regulation of lysosome trafficking machinery is altered in Batten disease might reveal novel regulatory defects in the affected neurons.

In the study by Uusi-Rauva *et al*. (2017)[28], the authors reprogrammed skin fibroblast from a CLN5 Batten disease patient to generate iPSCs and differentiated the iPSCs into neural cells. In the iPSC-derived neural cells with *CLN5* mutation, the authors revealed that LAMP1-stained organelles showed increased area and intensity. Although we did not find a change in lysosomal area and intensity with CLN5 inhibition in our iPSC-derived CLN5-deficient human neurons, we did detect an increased number of non-motile lysosomes (Figure 2I) in CLN5-deficient neurons with the overall number of lysosomes remaining the same (Figure 2E). The increased number of non-motile lysosomes explains the accumulation of lysosomes, seen in the CLN5-deficient neurons (Figure 2A, 2^nd^, 3^rd^ and 4^th^ panels), and as reported by Uusi-Rauva *et al*. Another study of CLN5 in HeLa cells by Yasa *et al*. (2021)[19] suggested that CLN5^KO^ HeLa cells were not able to move lysosomes as efficiently as wild type HeLa cells. In our study, impaired anterograde movement in the CLN5-deficient human neurons could indicate that the neuronal projections are being exempt from getting their waste cleared, which could cause neuronal toxicity at the synapses. Interestingly, we also observed that the increased non-motile lysosomes seemed to be accumulated in the cell body of the CLN5-deficient neurons (Figure 2A), which suggests that the lysosomes are stuck in the cell body and can’t move towards the projections, a phenotype supported by the Video S1 B, C. We also tested velocity of the lysosomes in the control and CLN5-deficient neurons, with no significant differences observed in anterograde or retrograde lysosomal velocity (Figure S1 A, B).

Overall, our iPSC-derived human neurons offer a wide range of applications from CRISPR manipulation to imaging specific organelle localisation and movement. Our human neuronal system offers an unprecedented platform for mimicking all forms of Batten disease in human neurons to study the interaction between the thirteen *CLN* genes in the development and progression of the disease, test neuronal phenotypes as well as screen therapeutic candidates for Batten disease.

---

**Figure S1.**
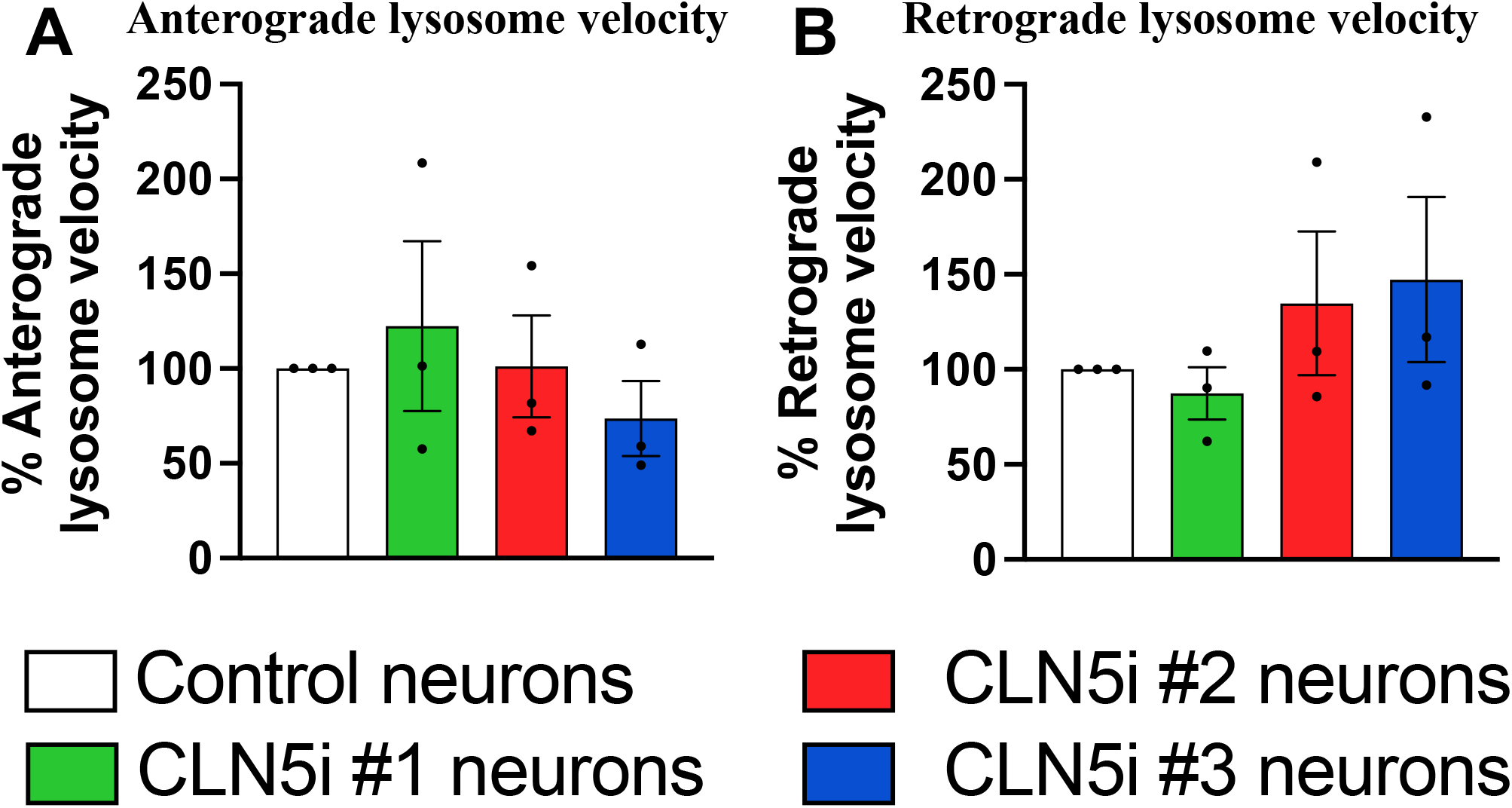
CLN5 deficient human neurons did not show a difference in the velocity of the motile lysosomes. CLN5-deficient neurons did not show any difference in the anterograde (A) and retrograde (B) velocity of lysosomes compared to the control neurons. The velocities of the lysosomes were measured using kymograph analysis as described in *Section 2.3*.

**Video S1: CLN5-deficient human neurons showed aberrant lysosomal movement**. Time lapse videos of acidic lysosomes were obtained for 5 minutes on a confocal microscope. CLN5i neurons (B, C) showed aberrant movement of the lysosomes (see encircled lysosomes), i.e., the lysosomes seem to be stuck and lacking a uni-directional movement as compared to the control neurons (A). Scale bar 20 μm.

## Author Contributions

Conceptualization, I.B., M.E.W., S.M.H..; methodology, I.B. and R.H.; validation, I.B.; formal analysis, I.B.; investigation, I.B., S.M.H. resources, M.E.W., S.M.H.; data curation, I.B.; writing–original draft preparation, I.B.; writing–review and editing, I.B., S.M.H.; supervision, S.M.H..; project administration, I.B., S.M.H.; funding acquisition, S.M.H. All authors have read and agreed to the published version of the manuscript.

## Institutional Review Board Statement

The study was conducted according to the guidelines of the Declaration of Helsinki and approved by the University of Otago Ethics Committee (application code: APP201858, approval number: GMC100228 and date of approval: 2^nd^ March 2015) for the import and use of genetically modified iPSCs. The iPSCs were obtained from Dr. Michael Ward at the National Institute of Health, USA.

## Acknowledgments

This study was made possible by the funding from the Neurological Foundation of New Zealand. The authors thank several group members of the Hughes laboratory, namely, Dr. Lucia Schweitzer, Ms. Jennifer Palmer, and Ms. Kirstin McDonald for their intellectual input towards the development of the manuscript. The authors thank the Otago Micro and Nanoscale Imaging (OMNI) facilities for the access to the flow cytometry and confocal microscopy. We thank Jonathan Weissman and Dider Trono for their generous gifts of the Addgene plasmids #60955, #12259 and 12260.

