## Supplementary Table for "Deficiency of the lysosomal protein CLN5 alters lysosomal function and movement"

### Table S1: Primary antibodies used

| Target protein | Species | Mono/polyclonal | Dilution used | Vendor | Catalogue no. |
|---|---|---|---|---|---|
| MAP2 | Guinea pig | Polyclonal | 1:2000 | Synaptic systems | 188004 |
| HOMER 1B/C | Rabbit | Polyclonal | 1:1000 | Synaptic systems | 160022 |
| SYNAPTOPHYSIN | Mouse | Monoclonal | 1:1000 | Synaptic systems | 101011 |
| TAU | Guinea pig | Polyclonal | 1:2000 | Synaptic systems | 314004 |
| ANKYRIN G | Mouse | Monoclonal | 1:200 | Invitrogen | 338800 |
| CLN5 | Rabbit | Monoclonal | 1:1000 | Abcam | 170899 |
| ACTIN | Mouse | Monoclonal | 1:2000 | Novus Biologicals | NB100-74340 |

### Table S2: Secondary antibodies used

| Target protein | Species | Dilution used | Vendor | Catalogue no. |
|---|---|---|---|---|
| *Anti Guinea Pig IgG (H+L) Highly Cross-Adsorbed Secondary Antibody, Alexa Fluor 647* | Goat | 1:1000 | Invitrogen | A-21450 |
| Anti-Rabbit IgG (H+L) Highly Cross-Adsorbed Secondary Antibody, Alexa Fluor 488 | Goat | 1:1000 | Invitrogen | A11034 |
| *Anti-Mouse IgG (H+L) Highly Cross-Adsorbed Secondary Antibody, Alexa Fluor 594* | Goat | 1:1000 | Invitrogen | A11032 |
| Anti-Mouse IgG (H+L) Highly Cross-Adsorbed Secondary Antibody, Alexa Fluor 488 | Goat | 1:1000 | Invitrogen | A11029 |
| IRDye® 680RD Anti-Mouse IgG Secondary Antibody | Goat | 1:20000 | Licor | 926-68070 |
| IRDye® 800RD Anti-Rabbit IgG Secondary Antibody | Goat | 1:20000 | Licor | 926-32211 |

### Table S3: qPCR probes used

| Target protein | Unique assay ID | Dilution used | Vendor | Catalogue no. |
|---|---|---|---|---|
| *CLN5* | qHsaCEP0050205 | 1:20 | Bio-Rad | 10031228 |
| *GAPDH* | qHsaCEP0041396 | 1:20 | Bio-Rad | 10031226 |
